# Trophic Efficiency Facilitates Larval Shortbelly Rockfish (*Sebastes jordani*) Development

**DOI:** 10.1101/2023.10.18.562988

**Authors:** Garfield T. Kwan, Kamran A. Walsh, Andrew R. Thompson, Noah J. Ben-Aderet, H. William Fennie, Brice X. Semmens, Rasmus Swalethorp

## Abstract

Identifying the factors influencing fish recruitment is critical for fishery management, and failure to do so can have major ecological and economic consequences. Many hypotheses over the past century have been proposed, and the recently postulated Trophic Efficiency in Early Life (TEEL) hypothesis argues that a shorter food chain length can result in more efficient energy transfer from primary producers to young fishes, thereby increasing growth rate and larval condition, reducing early-life mortality and ultimately leading to a stronger recruitment cohort. To test this hypothesis, we analyzed the trophic position (TP) through compound-specific isotopic analysis of amino acids, as well as otolith microstructure and stomach content of larval shortbelly rockfish (*Sebastes jordani*). Results show larval rockfish that ate lower TP prey were both heavier and faster growing. This suggests the trophic characteristics of early life diet are critical to larval survival, and provide evidence in support of the TEEL hypothesis.

## Introduction

Recruitment (the survival of larvae to juvenile or adult) is likely the single most important driver of adult population sizes for highly fecund fishes [1,2]. This dynamic is especially prevalent in short-lived marine fishes, which can fluctuate inter-annually by many orders of magnitude [3] and has major economic and ecological implications. For instance, Peruvian anchovetta (*Engraulis ringens*) in the Humboldt Current accounts for 73% of 2019 fishery landings valued at $3.5 billion USD [4]. They have undergone several booms and busts and their revenue fluctuated by several orders of magnitude over the last ∼150 years [5]. In the California Current Ecosystem (CCE), fluctuations in Pacific sardine (*Sardinops sagax*) and northern anchovy (*Engraulis mordax*) abundance are linked to variability in pinniped pup [6] and sea bird conditions [7]. Meanwhile, rockfish (*Sebastes* spp.) fisheries in California have historically exceeded $1 billion USD [8,9], but intense commercial and recreational fishing pressure dramatically reduced their numbers. Subsequent implementation of management plans and marine protected areas, coupled with favorable recruitment conditions, appeared to help rebuild certain rockfish populations [10]. However, to improve predictions and management strategies there is a great need to identify the factors that influence early life success and fish recruitment that drives population dynamics.

Hypotheses over the past century sought to explain the causes of recruitment variability, and they can be condensed to a single question: are larvae able to feed on prey that will allow them to grow faster and minimize predation? [11–15]. Past hypotheses focused on quantity of prey ingested since greater prey abundance generally leads to faster growth rate [16], however larval fishes are not simply generalist feeders [17]. Recently, the Trophic Efficiency in Early Life (TEEL) hypothesis postulates shorter food chain length through an “optimal” prey results in more efficient energy transfer from primary producers to young fishes, thereby increasing larval survival and ultimately leading to a strong recruitment cohort [18]. The TEEL hypothesis appears to explain most of northern anchovy (*Engraulis mordax*) population variability over a 45-year period, and food chain length/trophic position (TP) of their prey was quantified using high precision Compound Specific Isotopic Analysis of Amino Acids (CSIA-AA) technique. Here, we provide the first test of TEEL using larval shortbelly rockfish (*Sebastes jordani*).

Shortbelly rockfish are one of the most abundant rockfishes in the CCE, and they are an important forage species for many piscivorous fishes, sea birds, and marine mammals [19,20]. They grow to ∼35 cm, are reproductively active by 2 years and most live only to ∼12 years [19]. Unlike most rockfishes, shortbelly rockfishes are semi-pelagic and capable of explosive population growth fueled by extremely high annual recruitment [21]. Although not commercially fished, shortbelly rockfish can greatly impact fishery management. Shortbelly rockfish have expanded northward as populations exploded in recent years due to multiple high recruitment classes [22,23]. As a result many shortbelly rockfishes were taken as bycatch by the Pacific hake (*Merluccius productus*) fishery off the Oregon coast, which threatened to prematurely shut down the hake fishery [24]. This highlights the need to understand the mechanism(s) driving fishery recruitment dynamics, which remains a fundamental issue in fisheries science despite over a century of research [1,11]. In this study, we used a combination of CSIA-AA, otolith microstructure and gut content analysis to determine whether prey TP affects shortbelly rockfish larval condition in accordance with the TEEL hypothesis.

## Methods

### Sample collection

We used 95% ethanol-preserved larval rockfish samples collected by the California Cooperative Oceanic Fisheries Investigations (CalCOFI) program and archived at the Ichthyoplankton Collection curated by the NOAA Southwest Fisheries Science Center (La Jolla, CA, USA). *S. jordani* were identified via mitochondrial cytochrome *b* gene sequencing in a previous study [25]. To reduce variability, we selected larval *S. jordani* of comparable size (total length: 6.56 ± 0.23 mm; dry weight: 0.28 ± 0.03 mg; mean ± SEM) and collected during winter cruises at comparable CalCOFI sampling stations (Line(L)80 - Station(S)51, L80-S55, L83.3-S51, and L83.3-S55) during the years 2005, 2006, 2008, 2010, 2012, and 2013 (n=5 per year; Supplemental Figure 1). In total, 30 *S. jordani* were selected.

### Morphometric, Otolith, and Gut Analysis

*S. jordani* were placed on a microscope slide with deionized water, and imaged with a DSLR camera (Nikon D7000, Tokyo, Japan) under a compound microscope (Leica DMLB, Wetzlar, Germany) for standard length (SL) measurements. Next, sagittal otoliths were isolated and digestive organs were dissected for respective analyses. The remain portion of each larval *S. jordani* were frozen at -80°C for 24 hours, freeze-dried for 24 hours, then weighed on a microscale for dry weight. Samples were returned to the -80°C until CSIA-AA processing.

Dissected otoliths were mounted onto slides using clear nail polish, then Z-stacked imaged and digitally focus-stacked (Helicon Focus, version 8.0.4) with a compound microscope (1000x; Leica DMLB) and DSLR camera (Nikon D7000). Images were analyzed in R [26] using the package *RFishBC* [27] to obtain otolith core (center to post-rostral edge of extrusion check), age and increment widths (number and width of daily rings beginning at the first growth increment, respectively). Unfortunately, 7 sets of otoliths were lost during the dissection process.

Digestive tracts were dissected at 50x magnification, and stomach contents were identified, photographed, and measured with an eyepiece micrometer to the nearest 0.2μm. Recovered prey were categorized based on taxonomic grouping, growth stage, average length, average width, average number found within rockfish larvae with non-empty guts, percent numerical contribution to diet, average carbon weight, and percent carbon biomass contribution to diet. For copepodites length was measured as prosome length while total length was measured for all other prey groups. Carbon weights (μg) were estimated using conversion factors from existing literature (Supplemental Table 1).

### CSIA-AA and Trophic Position Estimation

This protocol follows those detailed in Swalethorp *et al*. [28]. Larvae were hydrolyzed in0.5 ml of 6N HCl for 24 h at 90°C. Hydrolyzates were then dried on a Labconco centrifugal evaporator under vacuum at 60°C. Particulates were removed by re-dissolving the hydrolyzates in 0.5 ml 0.1N HCl and filtering through an IC Nillix – LG 0.2-μm hydrophilic PTFE filter. Samples were re-dried before re-dissolving in 100 μl of 0.1% trifluoroacetic acid (TFA) in Milli-Q water, and stored at -80°C until AA separation. We used an Agilent 1200 series High Performance Liquid Chromatography (HPLC) system equipped with degasser (G1322A), quaternary pump (G1311A), autosampler (G1367B), and Realtek fixed flow splitter (5:1) which directed the flow to an analytical fraction collector (G1364C) and an Evaporative Light Scattering Detector (385-ELSD, G4261A), respectively. Amino acids (AA) were separated ona reverse-phase semi-preparative scale column (Primesep A, 10 × 250 mm, 100 Å pore size, 5 μm particle size, SiELC Technologies Ltd.) using a 120 minutes ramp solvent program 0.1% TFA in Milli-Q water (aqueous phase) and HPLC grade acetonitrile (ACN, organic phase). The fraction collector was programmed to collect glutamic acid (Glu) and phenylalanine (Phe) in 7 ml glass tubes based on elution time. Collection quality was assessed by comparing chromatograms with set collection times, and only when ≥ 99% of the peak areas fit within the collection windows where they accepted. Whole larval samples were injected onto the column to collect enough AA material for nitrogen (N) isotope analysis (≥1 μg N). Collected AAs were dried in the centrifugal evaporator at 60°C, dissolved in 40 μl of 0.1N HCl, and transferred to tin capsules (Costech, 3.5x5 mm). Capsules were then dried overnight in a desiccator under vacuum.

AA nitrogen isotopic analyses were performed at the Stable Isotope Laboratory at the University of California, Santa Cruz (UCSC-SIL) on a Nano-EA-IRMS system designed for small sample sizes (0.8-20 μg N). The automated system is composed of a Carlo Erba CHNS-O EA1108 Elemental Analyzer connected to a Thermo Finnigan Delta Plus XP Isotope Ratio Mass Spectrometer via a Thermo Finnigan Gasbench II with a nitrogen trapping system. δ^15^N values were corrected for size effects and instrument drift using Indiana University acetanilide, USGS41 Glu and Phe standards and correction protocols (see https://es.ucsc.edu/#silab) based on procedures outlined by Fry *et al*. [29].

N isotopic composition was measured consistently in 21 of 30 samples, with the rest lost during processing or discarded due to insufficient mass. δ^15^N of Glu and Phe are not significantly affected by ethanol preservation [28]. Trophic positions (TP) were calculated using and β and TDF values from Bradley *et al*. [30] and equation 1 (Supplemental Table 3).

### Statistical Analysis

All analysis was performed using R (version 4.0.3). Age and TP effects on SL and mass of larval *S. jordani* were analyzed with Bayesian hierarchical models encoded with the R package *‘rethinking’* [31]. Four Hamiltonian Markov chains with 1000 iterations were used. Bayesian inference is considered to lend advantages over frequentist methods when analyzing small sample sizes due to the inclusion of prior information, which increases robustness of analyses by incorporating current knowledge of expected outcomes into the model [32]. Since all variables were standardized, a mean of 0 and standard deviation of 1 were included as informative priors for each predictor variable. Normally distributed posterior distributions were summarized using central tendencies and variance, in this case the mean and 89% compatibility interval of each predictor variable. Sample station was included as a random effect (*varying intercept*) in both SL and body weight models, which were fit to the equation 2-9 (Supplemental Table 3). Posterior predictive checks were used to examine the predictive accuracy of the models, and model convergence was confirmed graphically using traceplots (Supplemental Figure 2).

To test the effects of TP on larval growth, another Bayesian hierarchical model in *‘rethinking’* was fit to the data with increment widths as the dependent variable and TP and increment number as independent variables. The interactive effect of increment number and TP was also included as an independent variable, with individual larvae included as a random intercept as denoted in equations 10-17 (Supplemental Table 3).

## Results and Discussion

As expected, age strongly influenced larval length and weight, indicating older larval *S. jordani* were both longer and heavier than their younger counterparts (Figure 1). In addition, feeding on prey at lower TP did not affect larval length (Figure 1A), but strongly increased larval weight (Figure 1B). In general, the condition of a fish corresponds to its weight at a given length, and forms the premise of which Fulton’s condition factor *k* is built upon [33]. Moreover, rationing experiments in adult [34] and larval fishes [35,36] indicate weight increase depends greatly on feeding conditions, whereas length can increase despite sub-optimal feeding conditions. This suggests the condition of *S. jordani* is related to the TP of larval prey, and by extension the efficiency by which energy is transferred from the base of the food chain up to and assimilated by the larvae [18].

**Figure 1.**
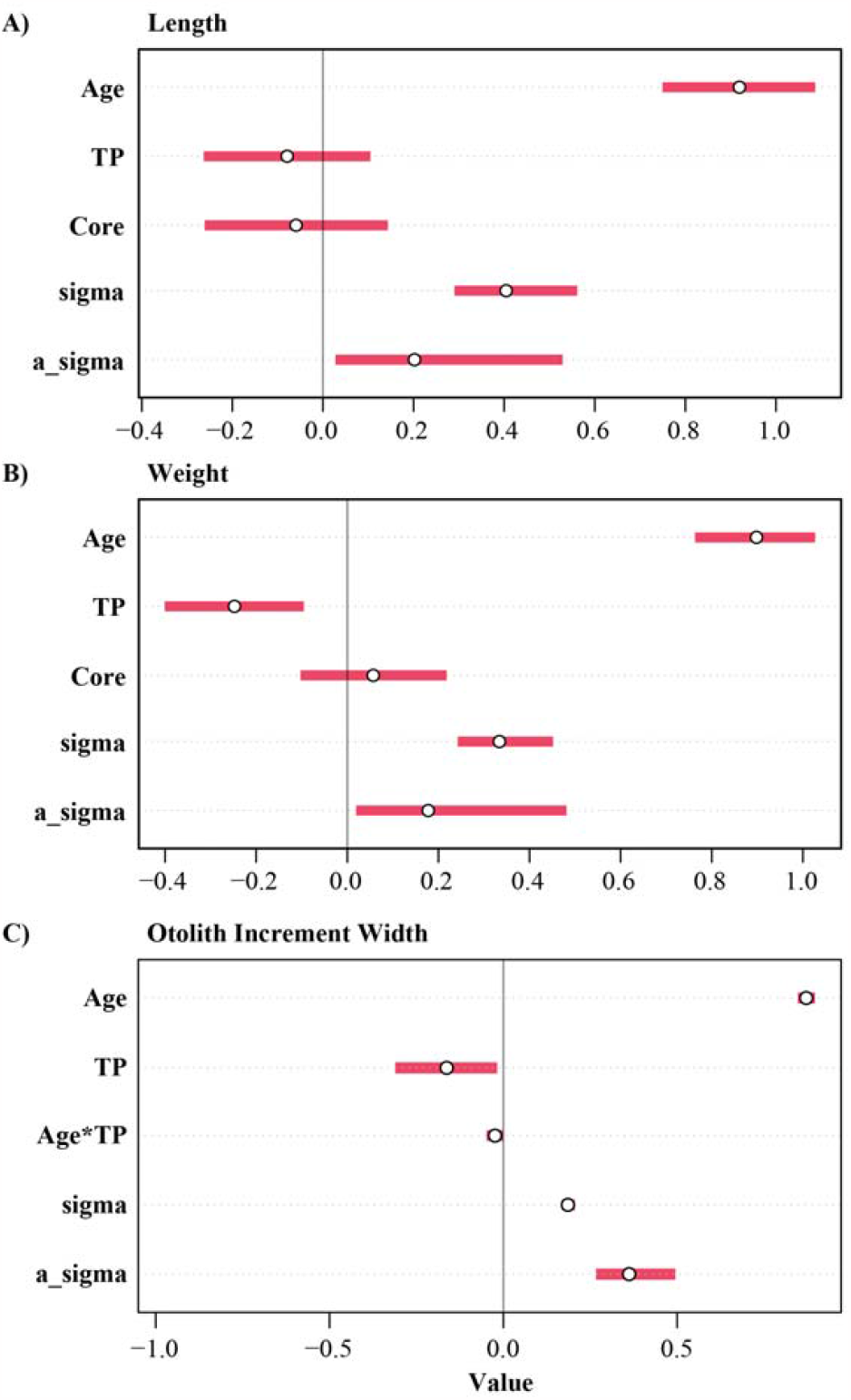
Predictors of larval *S. jordani* size and otolith increment width. **A)** Bayesian hierarchical model output indicates age is a strong predictor of *S. jordani* length, whereas **B)** both age and lower trophic position (TP) are strong predictors of *S. jordani* weight. In addition, Bayesian hierarchical model output indicates age, trophic position (TP), their interaction, and individual effects are all strong predictors of *S. jordani* otolith increment width. N=16. Data shows mean ± 89% compatibility interval. *sigma* represents the standard deviation of the normal distribution estimated by the model. *a_sigma* represents the variance of random effect.

Since TP integrates feeding history over the first days/weeks of life, we examined the otolith microstructures of larval *S. jordani* (6 to 25 days post extrusion) to assess whether trophic efficiency affects otolith size-at-age and growth rate. As expected, otolith increment width appears to strongly vary across individuals (Supplemental Figure 3). Even so, feeding on prey at lower TP strongly increased increment width, and this increase appear to magnify as the larvae age (Figure 1C). To visualize this, we parsed out larval *S. jordani* into a low and a high TP group (low TP: 1.99 ± 0.12; high TP: 2.75 ± 0.08; mean ± SEM; Figure 2). We found larval *S. jordani* feeding on prey at a low TP consistently exhibited faster otolith growth histories (Figure 2A) and larger otolith size-at-age (Figure 2B). Rapid growth has been identified as an important driver of larval survival: the stage duration hypothesis [37] argues mortality rate decreases as larvae develop and thus accelerated growth rate will decrease the duration fish spend in their vulnerable larval stage [38,39]. Empirically, slower growing individuals were found to be selectively preyed upon in a variety of fish studies including Japanese anchovy *Engraulis japonicus* [40] and quillback rockfish *Sebastes maliger* [41]. Our results imply that feeding on lower trophic level prey facilitates larger size-at-age and faster growth in larval *S. jordani*.

**Figure 2.**
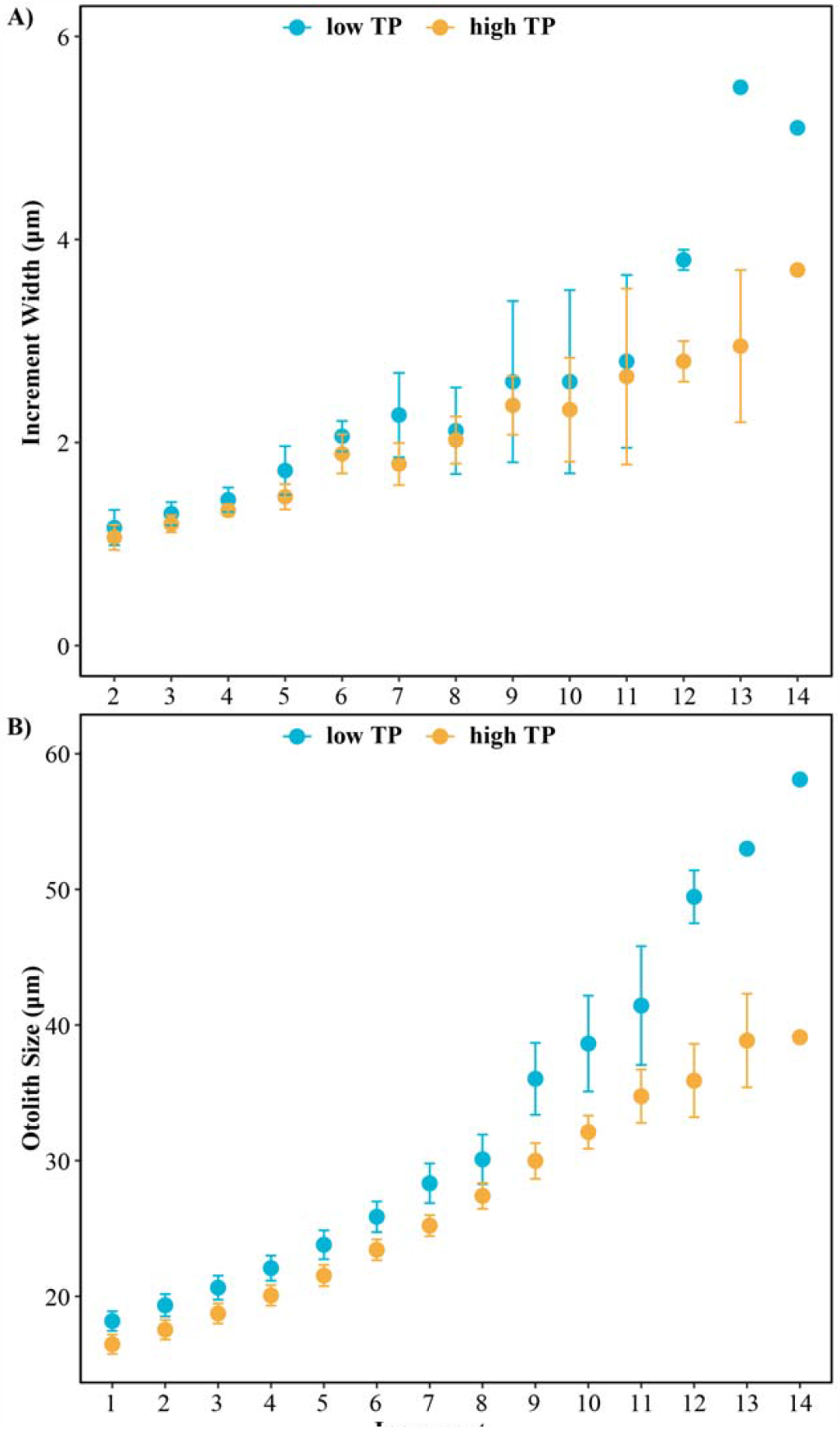
The relationship between increment width and otolith size against trophic position (TP). *S. jordani* with low TP (1.99±0.12) have **A)** faster otolith growth histories and **B)** larger otolith size-at-age compared to those with high TP (2.75±0.08). N=8 per treatment. Data shows mean ± SEM.

Larval size-at-age is postulated to positively impact survival as it influences swimming efficiency, prey capture and predator evasion [42,43]. In past studies, larger otolith core size (which reflects larval size at hatch/extrusion) was shown to be positively correlated with survival [38,39,44], and the effect magnified with age (45). Here, however, otolith core size did not affect larval *S. jordani* length or weight. Potential reasons for the lack of effect may be a small sample size or because any advantages related to larger core size would be inconsequential if there were unsuitable and/or insufficient prey available in the environment. Altogether, further research is necessary to explore the interactive relationships between TP and otolith core size and their roles as indicators of early larval success.

Next, we sought to identify the low TP food source responsible for heavier and faster growing larval *S. jordani*. Larval fishes are not simply generalist feeders [17], and evidence suggests the consumption of specific prey taxa can boost larval growth and survival [45–48]. Gut content analysis reveals the majority of the larval *S. jordani* diet is composed of *Calanoida* copepodites (41.73%), *Calanoida* nauplii (17.51%), and unidentified Copepoda copepodites (14.52%; Supplemental Table 2). Although larvae in the high TP group tend to consumed more carbon mass than the larvae in the low TP group (Supplemental Figure 4), their differences were not significant across total prey, *Calanoida* copepodite, and *Calanoida* nauplii (Supplemental Figure 4). This suggests larval *S. jordani* TP may be determined lower in the food chain, and not through prey switching by the larvae. Moreover, our inability to identify most prey items to species may have hindered our capacity to determine which prey optimizes energy transfer [2]. Further research (e.g. prey identification through metagenomics [49] is needed to understand the interplay between prey quality vs quantity, and what determines prey TP in larval fish diet.

Our results support the recent TEEL hypothesis [18]: larval feeding on prey at lower TP allowed for a more efficient energy transfer, resulting in faster growth and heavier size. We cannot rule out prey nutritional quality and larval food assimilation efficiency having some impact on the observed changes in TP [50–52]. Nevertheless, the end result is still less energy reaching the larvae lending support to TEEL, although indicates a need to further improve our mechanistic understanding of TEEL. While energy transfer appears to be an important component of larval survival, there are likely other factors (e.g. maternal investment, size-at-hatch/extrusion, and species-specific difference of prey) influencing larval recruitment that the present study could not adequately explore. As Reuben Lasker once pointed out, the key to understanding recruitment dynamics is to determine *what* limits recruitment and *when* in the life cycle this occurs [53]. Altogether, this study showcases how the specifics of trophic connections leading up to the prey of larval fish can impact larval success, and a greater understanding of their connectivity will allow for better predictions of fish recruitment with implications for conservation and fishery management.

## Acknowledgement

This project was supported by Scripps Institution of Oceanography (University of California San Diego) Small Grants Program. GTK was funded by the NSF Postdoctoral Research in Biology National Science Foundation (NSF) Postdoctoral Research Fellowship in Biology (Award #1907334) and by the Delta Stewardship Council Delta Science Program under Grant No. (21045). RS was funded by an NSF-RAPID Award (OCE-2053719) to BXS. The contents of this material do not necessarily reflect the views and policies of the Delta Stewardship Council, nor does mention of trade names or commercial products constitute endorsement or recommendation for use.

We thank the CalCOFI seagoing crew including, but not limited to, Amy Hays, Sue Manion, Dave Griffith and Bryan Overcash. We are grateful to Noelle Bowlin and Bill Watson for their help in sample identification, and Russ Vetter was invaluable for implementing the preservation of plankton in ethanol. We thank Dylan Gomes (NOAA SWFSC) for his helpful manuscript feedback, Stuart Sandin for the use of the compound microscope, Lihini Aluwihare for the use of the High-Performance Liquid Chromatography system, and Colin Carney at the Stable Isotope Lab at University of California Santa Cruz.

## Figures

**Supplemental Table 1.**
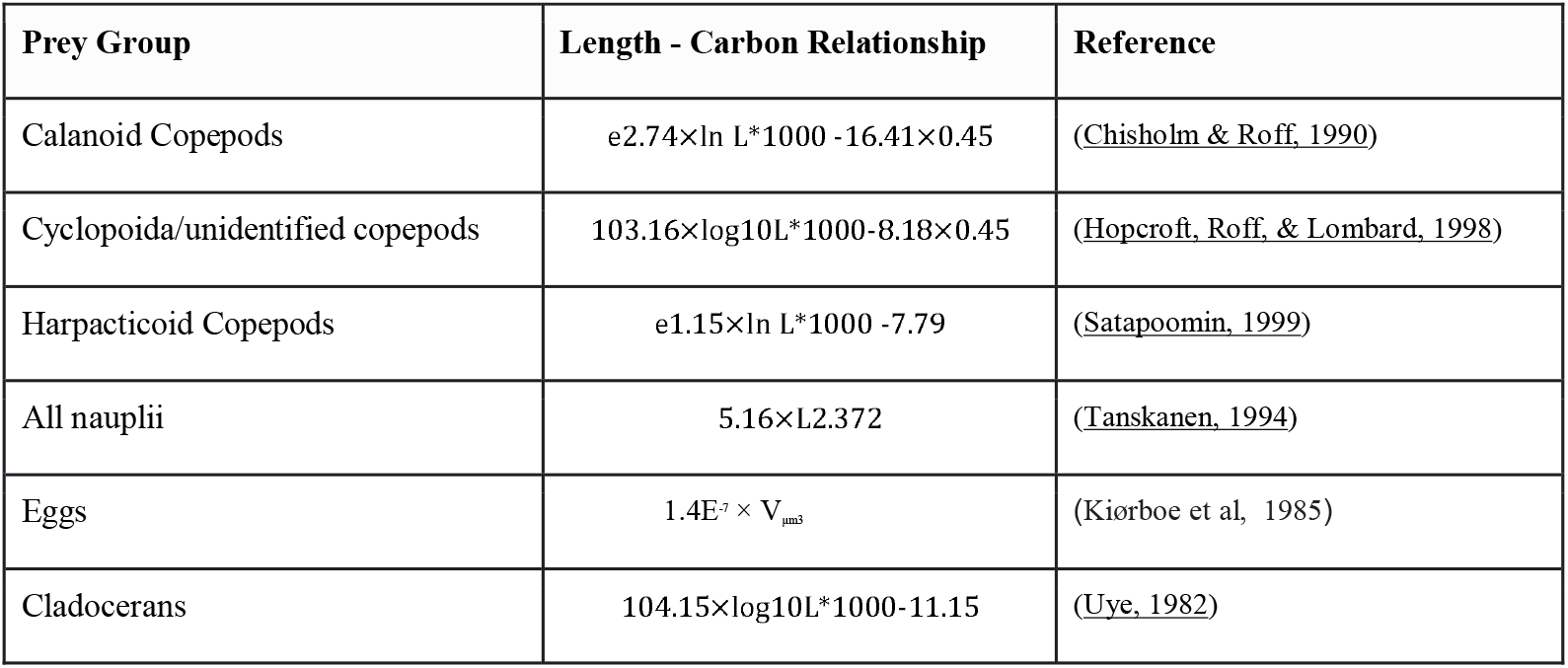
Length (mm) to carbon dry weight (μg) conversion factors for gut contents of *S. jordani*.

**Supplemental Table 2.** Diet metrics for recovered gut contents of *S. jordani*. Non-percentage values are presented as mean ± standard deviation.

| Prey | Average Length (µm) | Average Width (µm) | Average Number | Abundance (%) | Average C Weight (µg) | Biomass (%) |
| --- | --- | --- | --- | --- | --- | --- |
| *Calanoida Copepodites* | 431.58 ± 155.13 | 230.72 ± 88.23 | 2.92 ± 4.43 | 19.94% | 2.18 ± 3.94 | 41.73% |
| *Calanoida Nauplii* | 250.76 ± 68.15 | 141.96 ± 43.71 | 4.21 ± 4.20 | 28.77% | 0.92 ± 1.02 | 17.51% |
| *Copepoda Copepodites* | 395.11 ± 125.80 | 199.54 ± 42.23 | 1.17 ± 2.10 | 7.98% | 0.76 ± 1.45 | 14.52% |
| *Cyclopoida Nauplii* | 201.59 ± 44.97 | 107.39 ± 34.42 | 3.63 ± 3.85 | 24.79% | 0.45 ± 0.55 | 8.64% |
| Eggs | 186.68 ± 75.13 | 164.82 ± 29.20 | 0.46 ± 1.72 | 3.13% | 0.35 ± 1.45 | 6.63% |
| *Cyclopoida Copepodites* | 440.3 ± 100.88 | 212.75 ± 53.23 | 0.42 ± 1.10 | 2.85% | 0.33 ± 0.88 | 6.22% |
| *Other Nauplii* | 192.86 ± 53.14 | 124.88 ± 42.47 | 1.67 ± 2.10 | 11.40% | 0.19 ± 0.26 | 3.72% |
| Others | 282.13 ± 243.74 | 198.88 ± 177.04 | 0.17 ± 0.38 | 1.14% | 0.05 ± 0.19 | 1.02% |

**Supplemental Table 3.** Equations for the trophic position estimation and Bayesian hierarchical models used in this study. For equation 1, δ^15^N_Glu_ and δ^15^N_Phe_ are the isotopic compositions of the selected AAs. For equations 2-9, *y* is standard length (mm) or mass (mg), *r* is age (days), *w* is otolith core width (μm), and *t* is TP. For equations 10-17, *y* is otolith increment width (mm), *d* is increment number (day of life), and *t* is TP.

| Equation | Formula |
| --- | --- |
| 1 | $TP = \frac{\delta^{15}\text{N}_{\text{Glu}} - \delta^{15}\text{N}_{\text{Phe}} - \beta}{TDF_{\text{Glu-Phe}} + 1}$ |
| 2 | $y_i \sim \text{Normal}(\mu_i, \sigma_i)$ |
| 3 | $\mu_i = \alpha_{[\text{station}]} + \beta_r r_i + \beta_w w_i + \beta_t t_i$ |
| 4 | $\beta_r \sim \text{Normal}(0, 1)$ |
| 5 | $\beta_w \sim \text{Normal}(0, 1)$ |
| 6 | $\beta_t \sim \text{Normal}(0, 1)$ |
| 7 | $\alpha_{[\text{station}]} \sim \text{Normal}(0, \alpha_\sigma)$ |
| 8 | $\sigma_i \sim \text{Exp}(1)$ |
| 9 | $\alpha_\sigma \sim \text{Exp}(1)$ |
| 10 | $y_i \sim \text{Normal}(\mu_i, \sigma_i)$ |
| 11 | $\mu_i = \alpha_{[\text{individual}]} + \beta_t t_i + \beta_d d_i + \beta_{td} t_i d_i$ |
| 12 | $\beta_t \sim \text{Normal}(0, 1)$ |
| 13 | $\beta_d \sim \text{Normal}(0, 1)$ |
| 14 | $\beta_{td} \sim \text{Normal}(0, 1)$ |
| 15 | $\alpha_{[\text{individual}]} \sim \text{Normal}(0, \alpha_\sigma)$ |
| 16 | $\sigma_i \sim \text{Exp}(1)$ |
| 17 | $\alpha_\sigma \sim \text{Exp}(1)$ |

**Supplemental Figure 1.**
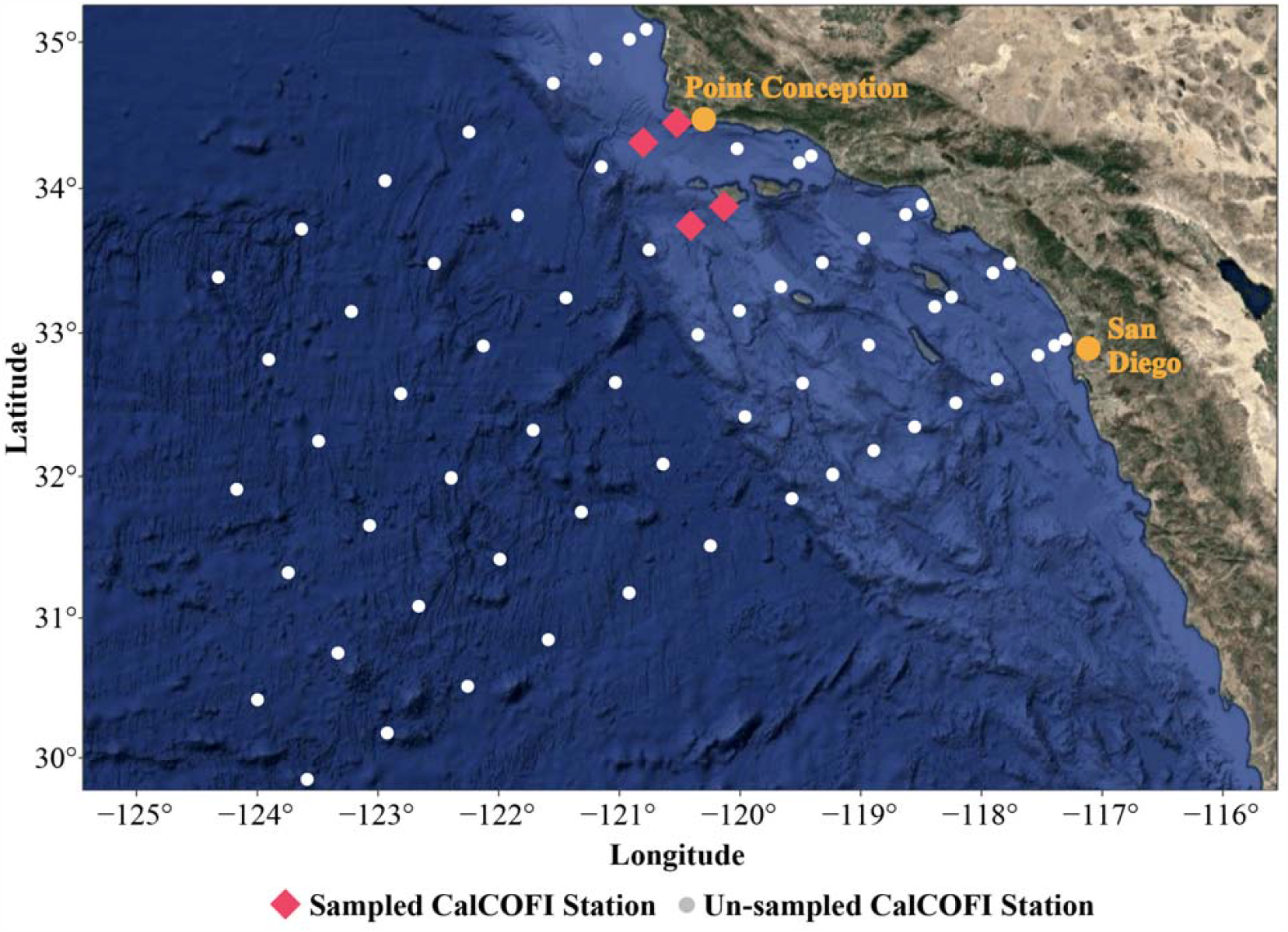
CalCOFI sampling stations and the sampling site for this experiment.

**Supplemental Figure 2.**
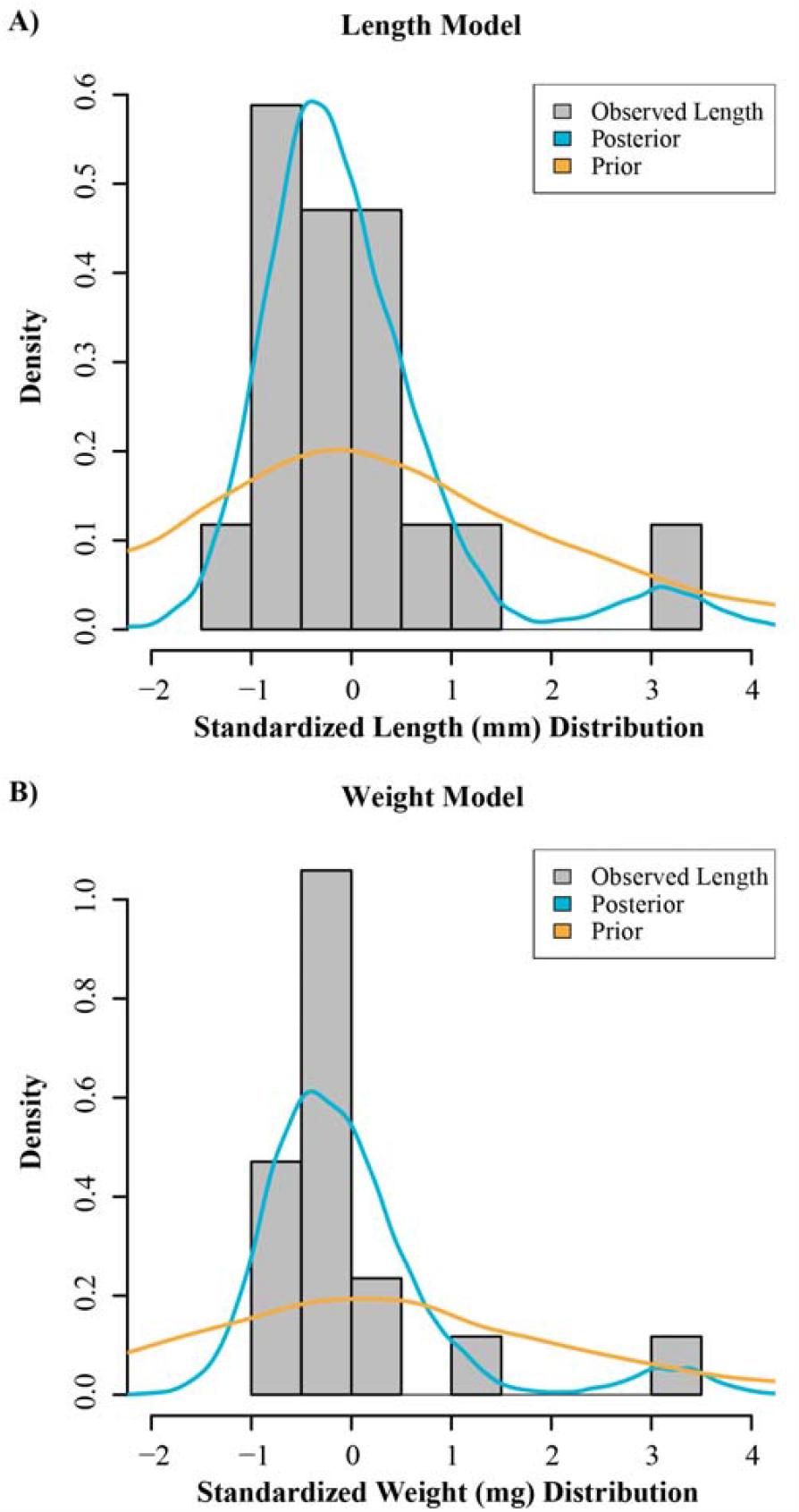
Posterior predictive checks for the **A)** TP to length and **B)** weight models.

**Supplemental Figure 3.**
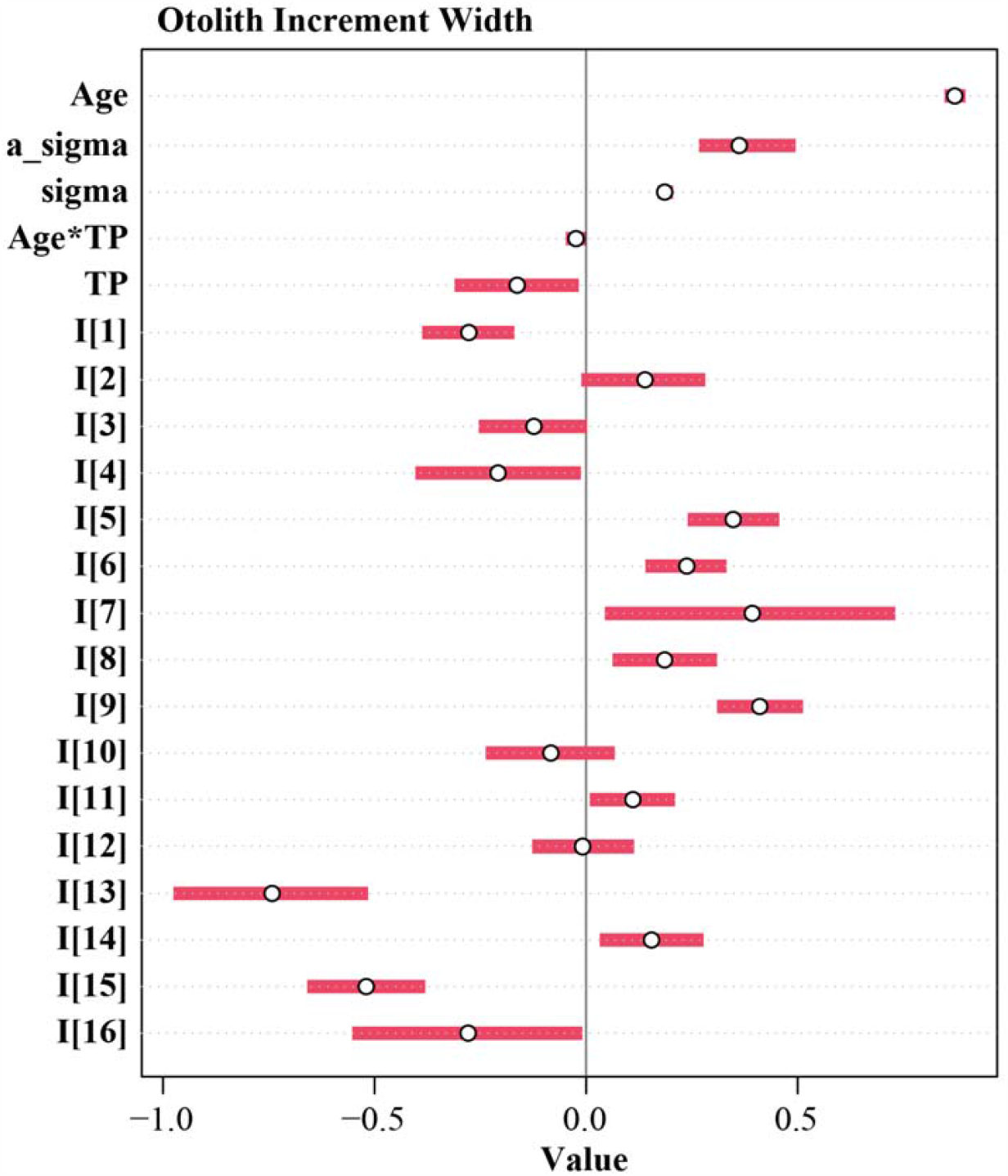
Predictors of larval *S. jordani* otolith increment width. Bayesian hierarchical model output predicting otolith increment width with all individuals (I) displayed alongside age, trophic position (TP), and their interaction. N=16. Data shows mean ± 89% compatibility interval. *sigma* represents the standard deviation of the normal distribution estimated by the model. *a_sigma* represents the variance of random effect. *I[X]* represents the random effect of individual larvae *X*.

**Supplemental Figure 4.**
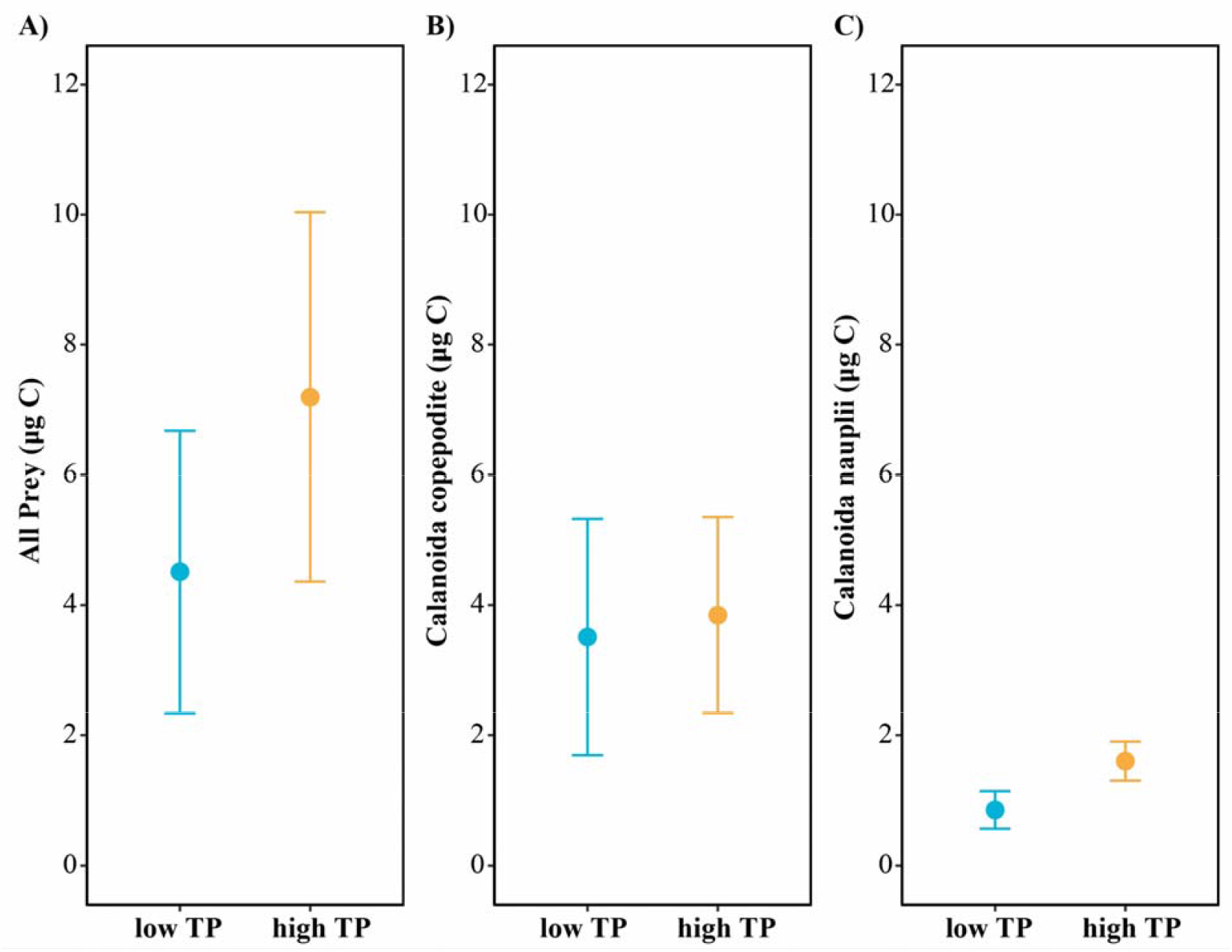
Comparison of carbon weight consumed across larval *S. jordani* from low and high larval trophic position (TP) groups. There was no significant difference in **A)** total prey (t = 0.7518, p-value = 0.4655), **B)** *Calanoida* copepodite (t = 0.0982, p-value = 0.9272), and **C)** *Calanoida* nauplii (t = 1.5603, p-value = 0.1498) consumed between larvae with low TP (1.99±0.12) and those with high TP (2.75±0.08). N=8 per treatment. Data shows mean ± SEM.

